# In vitro genotoxic and mutagenic potentials of combustion particles from marine fuels with different sulfur contents

**DOI:** 10.1101/2024.06.27.601016

**Authors:** Seongho Jeong, Jana Pantzke, Svenja Offer, Uwe Käfer, Jan Bendl, Mohammad Saraji-Bozorgzad, Anja Huber, Bernhard Michalke, Uwe Etzien, Gert Jakobi, Jürgen Orasche, Hendryk Czech, Christopher P. Rüger, Jürgen Schnelle-Kreis, Thorsten Streibel, Bert Buchholz, Thomas Adam, Martin Sklorz, Sebastiano Di Bucchianico, Ralf Zimmermann

**Affiliations:** Joint Mass Spectrometry Center (JMSC) at Comprehensive Molecular Analytics (CMA), Helmholtz Zentrum München, Ingolstädter Landstr. 1, 85764 Neuherberg, Germany; Joint Mass Spectrometry Center (JMSC) at Analytical Chemistry, Institute of Chemistry, University of Rostock, Albert-Einstein-Strasse 27, 18059 Rostock, Germany; University of the Bundeswehr Munich, Faculty for Mechanical Engineering, Institute of Chemical and Environmental Engineering, Werner-Heisenberg-Weg 39, 85577 Neubiberg, Germany; Research Unit Analytical BioGeoChemistry, Helmholtz Zentrum München, Ingolstädter Landstr. 1, 85764 Neuherberg, Germany; Chair of Piston Machines and Internal Combustion Engines, Faculty of Mechanical Engineering and Marine Technology, University of Rostock, Albert-Einstein-Strasse 2, 18059 Rostock, Germany

**Keywords:** Marine Gas Oil (MGO), Heavy Fuel Oil (HFO), Particulate Matter (PM), Ship emission, Genotoxicity, Mutagenicity

## Abstract

Ship emissions cause serious environmental impacts and adverse effects toward human health. Therefore, the International Maritime Organization (IMO) restricted the fuel sulfur content (FSC) of marine fuels: FSC must be <0.5% m/m or <0.1% m/m in sulfur emission control areas, covering a range of fuels from distillate diesel-like fuels to low-sulfur heavy fuel oils (HFOs). As a result, ship emissions, e.g., sulfur oxides and particulate matter (PM) have been reduced. However, how FSC correlates with the toxicological potential of ship emissions is still uncertain. The objective of this study was to understand how the physical and chemical properties of particulate emissions from a marine engine operating on five marine fuels with different FSCs influence their toxicological outcome. For this scope, cytotoxic, genotoxic, mutagenic, and pro-inflammatory potentials of collected particles were evaluated in lung cell model systems. The involvement of intracellular reactive oxygen species and xenobiotic metabolism was also explored. While PM from different fuels’ combustion resulted in up to approximately 20% of reduction of cytotoxicity at the highest concentration, other toxicological outcomes, including clonogenic and genotoxic potentials, showed a stronger trend with the polycyclic aromatic hydrocarbon contents in PM compared with FSC. This trend was supported by evidence of a significant increase in gene mutation frequency and alterations in cellular mechanisms induced by an aromatic-rich HFO with an intermediate FSC. In conclusion, apart from reducing FSC in marine fuels, additional particle abatement systems should be considered to reduce the adverse effects of particulate emissions from shipping operations on human health.

## Introduction

It is widely acknowledged that deteriorated air quality, characterized by elevated concentrations of airborne particulate matter (PM) and pollutants from combustion process, can significantly impact human health. Inhalation of such pollutants can lead to oxidative stress and inflammation (Delfino et al., 2011). These effects are associated with various deleterious health effects, including the development of cardiovascular disease (Lu et al., 2021) and respiratory diseases (Chauhan and Johnston, 2003). Among different emission sources, over the past few decades, human health and the environmental impact of ship emissions have become an increasingly urgent issue. Although regulations and advancements in after-treatment techniques have significantly reduced emissions from land vehicles (Eyring et al., 2005; Ramacher et al., 2020), ship emissions such as particulate matter and sulfur oxides (SO_x_) remain a significant concern. SO_x_ can have serious consequences causing acidification in water and soil (Hassellöv et al., 2013). Furthermore, it is one of the precursors for secondary particle formation in the atmosphere, accompanied by direct ship emission PM. As a result, these pollutants have gained attention as an important causative of morbidity and premature deaths, especially near harbors and in coastal cities (Corbett et al., 2007; Fuglestvedt et al., 2009; Hong et al., 2023).

In order to address the reduction of shipping emissions, the International Maritime Organization (IMO, 2008) adopted guidelines and defined fuel sulfur content (FSC) upper limits of 0.1% and 0.5% for ships operating inside and outside of sulfur emission control areas (SECAs), respectively. To meet IMO regulations, ships must be operated using low-sulfur fuels, such as marine gas oil (MGO), low-sulfur heavy fuel oil (HFO), or hybrid fuel. Alternatively, high-sulfur fuels can still be used if emission reduction technologies are installed to achieve sulfur emission levels similar to low-sulfur compliant fuels.

While it has been demonstrated that reducing the FSC can decrease particulate emissions from shipping operations (Moldanová et al., 2009; Sippula et al., 2014; Mueller et al., 2015), little is known about whether changing the type of marine fuel can significantly reduce the toxicological potential of emissions. Epidemiological evidence has estimated that implementing stricter SECA regulations could lead to health improvements (Brandt et al., 2013; Tang et al., 2020) due to reduction of emitted pollutants. While these estimates based on particle number and mass emissions have suggested that stricter SECA implication would cause positive effects on both human health and the environment, the impacts of PM-constituents derived from different fuel types on these areas remain poorly understood.

In the near future, the expected growth of global trade and international supply chains will potentially increase shipping emissions. Therefore, research that assesses their differential toxicological potentials is needed (Mueller et al., 2022). It has been shown that PM from HFO induced higher cytotoxicity and oxidative stress compared with that from Diesel oil (Wu et al., 2018). However, other studies have demonstrated that Diesel oil PM induced a broader range of responses in human lung cells, including energy metabolism, protein synthesis, and chromatin modification (Oeder et al., 2015; Sapcariu et al., 2016). To the best of the authors’ knowledge, no study has been conducted to investigate the effects of PM from MGO combustion on mammalian cells. Moreover, no *in vitro* assessments have been performed to evaluate the effects of PM generated from the combustion of HFOs with different compositions, including FSC and aromaticity.

This study investigates the toxicological effects of fine PM (PM_2.5_) ship emission on human alveolar lung cells, in which the deposition of fine PM dominates compared to other respiratory tracts (Cipoli et al., 2023). The particulate emissions are collected from combustion of different types of marine fuel combustion emissions on human lung cells. Specifically, this study addresses the scientific question if the sulfur content in different fuels is related to differences in biological responses to PM emission produced by the combustion of these fuels.

The objective of this study is to address the toxicological effects of PM from five different fuels’ combustion spanning an FSC range of 0.06% m/m to 2.4% m/m on lung cells with variations in concentration and exposure time. We analyzed the cytotoxic, clonogenic, genotoxic, and mutagenic potentials of the collected particles, as well as their oxidative reactivity, xenobiotic metabolism, and inflammatory properties in relation to physical and chemical parameters such as particle number size distribution, particle mass emission factors, organic compositions, and metal contents. By addressing these critical questions and concerns, this study aims to contribute to a deeper understanding of the toxicological effects associated with ship emissions and to provide valuable insights for policy development and mitigation strategies in the shipping industry.

## Material and Methods

### Characterization of the ship engine and marine fuel types

To mimic the realistic combustion processes of a marine engine with a stable and reproducible emission scenario, a four-stroke, single-cylinder, common rail research marine engine at the University of Rostock was used. The engine with a rated power of 80 kW at 1500 rpm was invented as a representative of a typical modern medium-speed marine engine and was equipped with a compressor-charged air-intake for fixable air-fuel equivalence ratios. More detailed specifications of the test engine can be found in previous studies (Mueller et al., 2015; Streibel et al., 2017). The engine was operated on fixed cycles at four different engine loads of 20, 40, 60, and 80 kW for each fuel type, corresponding to 25, 50, 75, and 100% of the maximum continuous rate, respectively. The duration and sequence of the engine loads were set according to the weighting factors of ISO 8178-4 E2, with a total cycle duration of 6 h. Specifically, the engine loads of 20 kW and 40 kW were maintained for 54 min, 60 kW for 180 min, and 80 kW for 72 min. The entire test cycle was run 3 times with each fuel type, followed by a change of the lubricating oil for the next test cycle with a different fuel type.

Five different marine fuels, namely, one MGO, and four HFOs, were used in our experiments. MGO is considered one of the highest quality marine fuels due to its low-sulfur and metal contents compared with HFO. It is regularly used in highly regulated areas such as the Baltic and North seas, as well as in coastal areas near North America (Fan and Gu, 2019). HFOs typically have much higher viscosity and density than MGO and contain residual fractions from refinery processes (Fritt-Rasmussen et al., 2018). Therefore, HFOs can contain a significant amount of metals (mainly Ni, V, and Fe), sulfur, and nitrogen. By definition, an HFO has a density greater than 0.9 g cm^-3^ at 15°C or a kinematic viscosity exceeding 180 mm^2^ s^-1^ at 50°C, allowing for a broad range of marine fuels from blending different product streams of the crude oil refining. For our study, MGO, low sulfur HFO (LS-HFO), and high-sulfur HFO (HS-HFO) were purchased as commercial marine fuels. Additionally, two blended noncommercial fuels, an ultra-low sulfur HFO (ULS-HFO_AR_) and a high sulfur HFO (HS-HFO_SYN_), containing high amount of aromatic compounds, including naphthalene, and phenanthrene derivatives, were investigated. Although the FSC is associated with the high boiling fractions of HFOs (Hsieh et al., 2013), LS-HFO showed the smallest mass loss in boiling distribution analysis until 400°C (Fig. S1). ULS-HFO_AR_ is highly similar to distillate fuels like MGO in terms of sulfur, and metal contents, viscosity, and flash point, despite its classification as an HFO and low combustion efficiency. This marine fuel refers to a clarified cycle oil, produced from fluid catalytic cracking and mainly contains alkylated 2- to 4-ring aromatics (Käfer et al., 2019). HS-HFO_SYN_ features typical properties of HFO but comprises distinct low- and high-boiling fractions in a ratio of approximately 30:70. The combustion efficiency of each fuel type is expressed in Table 1 as C/H ratio (m/m), calculated carbon aromaticity index (CCAI), and aromaticity index (AI) (Koch and Dittmar, 2006). The highest C/H, CCAI, and AI values of ULS-HFO_AR_ resulted in the lowest ignition quality, which led to a maximum engine load of 68 kW instead of 80 kW.

**Table 1.**
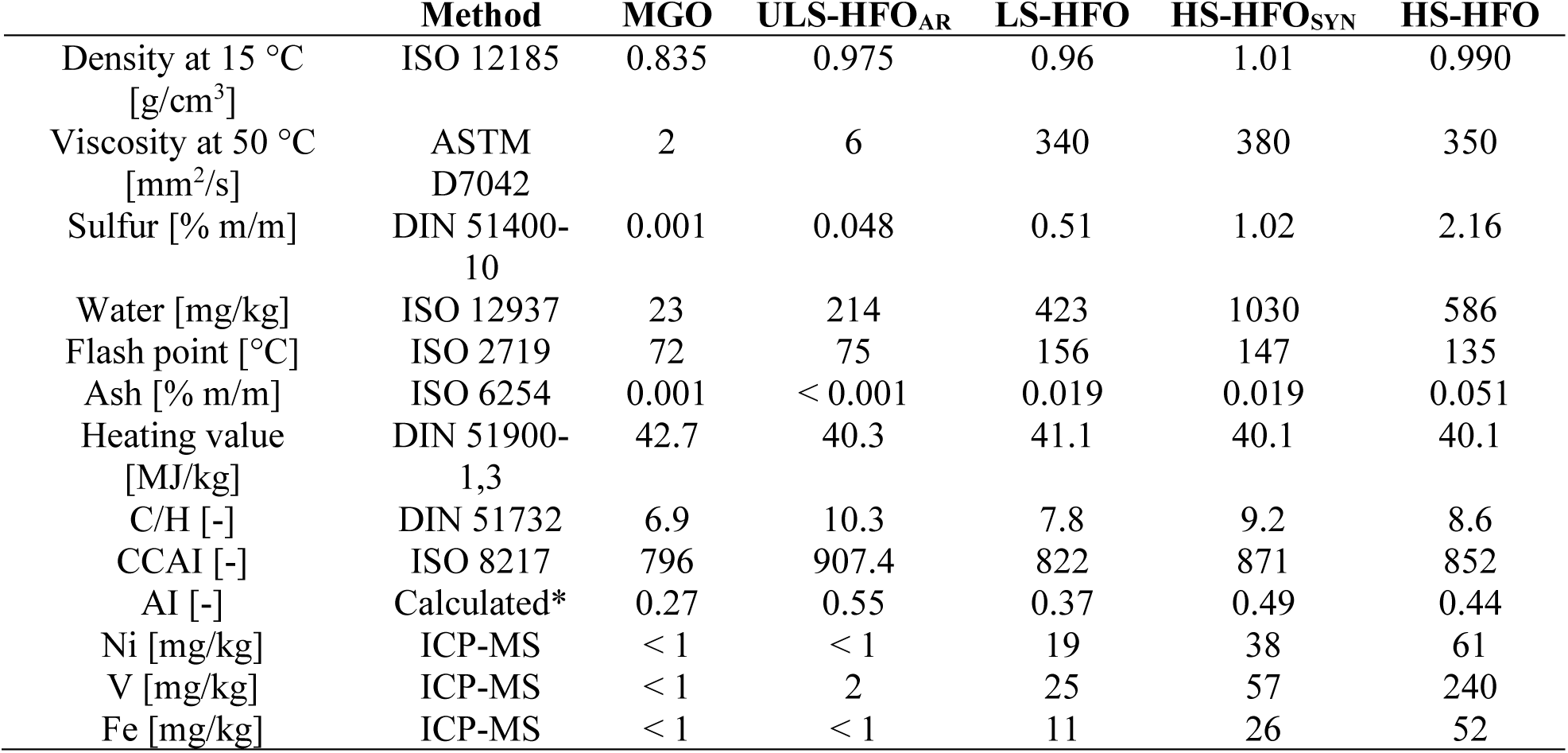
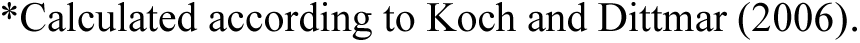
Physical and chemical properties of applied fuels.

|  | Method | MGO | ULS-HFO<sub>AR</sub> | LS-HFO | HS-HFO<sub>SYN</sub> | HS-HFO |
| --- | --- | --- | --- | --- | --- | --- |
| Density at $15^\circ\text{C}$<br>[g/cm<sup>3</sup>] | ISO 12185 | 0.835 | 0.975 | 0.96 | 1.01 | 0.990 |
| Viscosity at $50^\circ\text{C}$<br>[mm<sup>2</sup>/s] | ASTM<br>D7042 | 2 | 6 | 340 | 380 | 350 |
| Sulfur [% m/m] | DIN 51400-10 | 0.001 | 0.048 | 0.51 | 1.02 | 2.16 |
| Water [mg/kg] | ISO 12937 | 23 | 214 | 423 | 1030 | 586 |
| Flash point [$^\circ\text{C}$] | ISO 2719 | 72 | 75 | 156 | 147 | 135 |
| Ash [% m/m] | ISO 6254 | 0.001 | < 0.001 | 0.019 | 0.019 | 0.051 |
| Heating value<br>[MJ/kg] | DIN 51900-1,3 | 42.7 | 40.3 | 41.1 | 40.1 | 40.1 |
| C/H [-] | DIN 51732 | 6.9 | 10.3 | 7.8 | 9.2 | 8.6 |
| CCAI [-] | ISO 8217 | 796 | 907.4 | 822 | 871 | 852 |
| AI [-] | Calculated* | 0.27 | 0.55 | 0.37 | 0.49 | 0.44 |
| Ni [mg/kg] | ICP-MS | < 1 | < 1 | 19 | 38 | 61 |
| V [mg/kg] | ICP-MS | < 1 | 2 | 25 | 57 | 240 |
| Fe [mg/kg] | ICP-MS | < 1 | < 1 | 11 | 26 | 52 |

\*Calculated according to Koch and Dittmar (2006).

### Sampling setup and particle collection

The setup for particle collection and characterization is shown in Fig.1. Particle emissions from the engine exhaust were taken with two different dilution systems, one for the toxicological assessment and chemical characterization (offline) and the other for online particle physical characterization. To prevent vapor condensation on particle emissions, a 250°C heat transfer line was installed between the sampling point and dilution system for both sampling lines of offline and online instruments. The dilution system (Venacontra, Finland) for the toxicology study consisted of a porous tube diluter (PTD) in which clean compressed air flowed through pores in a cylindrical sampling volume to provide sheath flow and minimize the wall losses of vapors and particles. Then, the particles were guided into an ejector diluter (EJD), which thoroughly mixed the sample with the dilution air. To inhibit the condensation of sulfuric acid in the dilution system but to collect the particles with the highest yield, the dilution factor was set between 5 and 10, depending on the FSC, and its dew point at 25°C. CO_2_ probe GMP 343 (Vaisala, Finland) was utilized to calculate the exact dilution factor by continuously monitoring the CO_2_ concentration, comparing it with the raw gas and background CO_2_ concentration in dilution air, as previously described (Grigonyte et al., 2014). The coarse particles (PM_10_) from the exhaust emission were separated using a pre-cyclone and particles smaller than PM_10_ were collected in a commercially available sampling box (CGS GmbH, Germany), where a ceramic filter (1011756, CGS GmbH, Germany) was used to collect the particles directly on the filter surface. Particles were sampled with a volume flow of 50 L min^-1^ after appropriate dilution and cooling to 25°C. After the measurement cycle, the particles on the ceramic filter were carefully scratched off using a sterile stainless spatula and collected in a sterile glass vial for storage at −20°C.

**Fig. 1.**
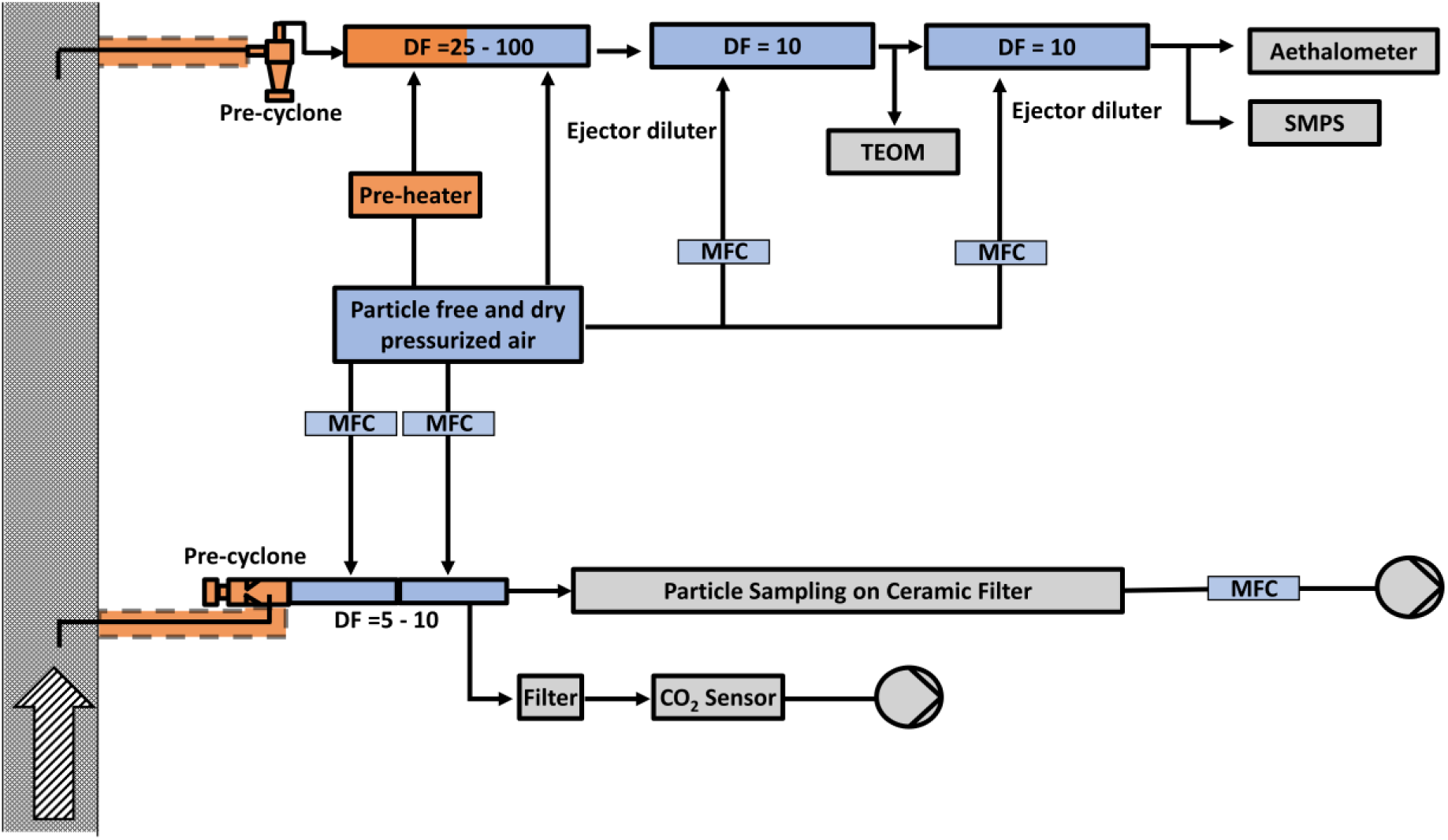
Scheme for the sampling setup. Arrow with stripe pattern indicates the ship emission from the engine. TEOM, tapered element oscillating microbalance; SMPS, scanning mobility particle sizer; DF, dilution factor; MFC, mass flow controller

To avoid any possible interruptions to the online equipment, a separate sampling point for particle characterization was established, and the exhaust was diluted using a two-stage EJD system (eDiluter, Dekati Ltd., Finland). The dilution ratio of the eDiluter was optimized on the basis of the detection limit of the online equipment, and the dilution factor was adjusted from 25 to 100, depending on the particle number concentration of different fuel types. Moreover, further dilution steps were established using EJDs (VKL 10, Palas GmbH, Germany) with a fixed dilution factor of 10.

### Online physical characterization

A comprehensive study of the particle size distribution, number, and mass concentrations was performed using a scanning mobility particle sizer (SMPS) and a tapered element oscillating microbalance (TEOM 1400a, Thermo Fisher Scientific, USA). The SMPS was operated at an aerosol flow of 0.3 L min^−1^ with an X-ray neutralizer (TSI, Model 3088, USA), an electrostatic classifier (TSI, Model 3082, USA), and a condensation particle counter (TSI, Model 3750, USA). The TEOM measured the particle mass concentrations at a sample flow rate of 3 L min^−1^, with a cap and inlet tube temperature of 50°C as the standard setting, according to Patashnick and Rupprecht (1991). To account for artificial changes in gaseous and particulate emissions during engine load changes, only measurements taken after a stabilization period of approximately 30 min were considered valid. Furthermore, an aethalometer (AE33-7, Magee Scientific, Aerosol, d.o.o., Slovenia) was used for real-time monitoring of the light absorption properties of PM to determine the equivalent mass concentration of black carbon (eBC).

### Polycyclic aromatic hydrocarbons (PAHs)

For the quantification of PM-bound PAHs, the collected PM was analyzed using direct thermal desorption comprehensive two-dimensional gas chromatography coupled to high-resolution time-of-flight mass spectrometry (DTD-GC × GC-HRTOFMS). The applied analytical method is based on previous studies (Schnelle-Kreis et al., 2005; Orasche et al., 2011) and adapted for the application on a Leco Pegasus HRT 4D system (Leco, USA) with an Optic 4 programmable vaporizing injector (GL Science, the Netherlands). Defined aliquots of approximately 1 mg of particles were homogenized with baked out, grounded sodium sulfate (1:1,000, w/w) which served as inert matrix for dilution. This approach facilitates subsequent sample handling steps and prevents overloading of the analytical setup. From these mixtures, precisely weighted amounts in the range between 20 and 40 mg were transferred to a GC-liner and spiked with an internal standard mixture, containing deuterated PAHs. Subsequently, the prepared samples were analyzed using DTD-GC × GC-HRTOFMS with three repetitions for each sample. Detailed information about thermal desorption, chromatographic, and mass spectrometric parameters can be found in Table S1. Quantification of 12 target PAHs was performed by internal standard correction and external calibration with the same approach as previously described (Schnelle-Kreis et al., 2005; Orasche et al., 2011).

### PM elemental content by ICP-OES

To perform elemental analysis, collected PM was suspended in 1 mL of sub-boiling distilled 65% nitric acid (Roth, Germany) overnight and subsequently diluted with 3 mL of Milli-Q water. The resulting PM suspension was filtered using one-way syringe filters (0.22 µm, VWR, Germany) and then analyzed using ICP-OES (Inductively coupled plasma atomic emission spectroscopy) (ARCOS, Germany). The results are demonstrated in µg mg^-1^ of particle mass, and the limit of quantification is shown in Tables 4 and S3. To determine the measurement uncertainty (*Unc)* for each element *(i),* equation 1 was used:

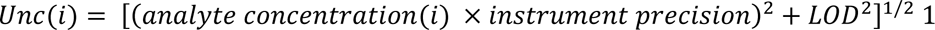

with LOD being the limit of detection.

### Mutagenic equivalent (BaPMEQ)

The mutagenic potency of collected PM was evaluated by calculating benzo[a]pyrene (BaP) mutagenic equivalent values (BaPMEQ, ng m^−3^), which are based on the mutagenic equivalency factors (MEF) relative to BaP (Durant et al., 1996). The mutagenic equivalents of sampled PM were calculated by multiplying the MEF by the concentration of each PAH (Table S2).

### PM suspension preparation

To prepare PM suspensions for *in vitro* testing, 0.05% w/v bovine serum albumin (BSA, CAS-9048-46-8, Fraction V, BioMol GmbH, Germany) water solutions were freshly prepared and diluted in cell culture medium according to the NANOGENOTOX protocol with minor modifications (Zijno et al., 2020). Briefly, 1% w/v solution of the BSA powder in ultrapure water was prepared, filtered using a sterile syringe filter with a pore size of 0.22 µm (16532-GUK, Sartorius, Germany), and further diluted to a concentration of 0.05% w/v in ultrapure water. Each type of PM was weighed using a microbalance (MC-1 AC210S, Sartorius, Germany) and aliquoted in a sterile glass vial (548-0509, VWR, Germany). As a prewetting procedure, 0.5% v/v of ethanol (Panreac AppliChem, Germany) was added to each PM sample in the glass vial. After 1 min of the prewetting procedure, the PM was suspended in 0.05% w/v BSA–water to a concentration of 2 mg mL^−1^ and sonicated (PALSONIC, ALLPAX, Germany) for 30 min in an icy water bath to prevent excessive heating while swirling and agitating every 5 min. The resulting PM suspensions were then diluted with high-glucose Dulbecco’s Modified Eagle Medium /Nutrient Mixture F-12 (CAS 31331-028, DMEM/F12, Gibco) to reach a PM concentration of 1 mg mL^−1^ and sonicated for an additional 5 min. After the second sonication step, the stock solutions were further diluted in the cell culture medium to achieve the desired PM concentrations.

### Physical characterization of PM suspension

To understand the physical properties of the PM in the exposure medium for the submerged experiments, the hydrodynamic particle diameter, the polydispersity index (PDI), and the zeta potential were measured. A well-dispersed PM suspension with a concentration of 50 µg mL^-1^ was freshly prepared for the PM from each type of fuel. The hydrodynamic size and PDI were measured using a micro cuvette (ZEN0040, Malvern Instrument Ltd., UK) at room temperature, with a backscatter angle of 173° (Nano ZSP, Malvern Instrument Ltd., UK). For zeta potential measurements of PM in DMEM/F12 cell culture media, the PM suspension was added to a folded capillary zeta cell (DTS1070, Malvern Instrument Ltd., UK). Each physical parameter was measured 10 times after a 120-s equilibration time for each sample.

### Cell culture and treatments

A549 human alveolar epithelial cells (CCL-185, ATCC^®^, USA) were routinely cultured and maintained in DMEM/F12 supplemented with 5% v/v heat-inactivated fetal bovine serum (FBS, 10500-064, Thermo Fisher Scientific, USA), 100 U mL^-1^ penicillin, and 100 µg mL^−1^ streptomycin (P4333, Sigma-Aldrich, USA) in a humidified incubator at 37°C and 5% CO_2_. For the exposure experiments, A549 cells (passages from 5 to 25) were seeded at a concentration of 7.6 × 10^4^ cells cm^−2^ on 24-well plates (Corning^®^, USA). After 24 h, PM dispersions were added to the cell cultures to a final concentration of 3.5, 15, and 55 µg cm^-2^, and cells were further incubated for 4 or 24 h. A549 cells cultured in DMEM/F12 were depicted as negative controls, and cells treated with 0.05 % w/v BSA–water in DMEM/F12 media were used as solvent control in all toxicological assays.

### Cell viability

Cell viability was evaluated using trypan blue exclusion assay. After 4 or 24 h exposure, A549 cells were washed twice with prewarmed PBS and detached using 0.05% trypsin–EDTA (T4174, Sigma-Aldrich, USA). Afterwards, 20 µL of the cell suspension was mixed with an equal volume of 0.4% w/v trypan blue solution (T8154-20ML, Sigma-Aldrich, USA), and the cell viability (percentage of unstained cells) was determined using a hemocytometer (Neubauer, Germany).

### Colony forming efficiency assay

Cell proliferation was investigated by measuring the ability of single cells to form a colony consisting of at least 50 cells and was demonstrated as clonogenicity at a given time point, which is closely related to cytotoxicity (Franken et al., 2006; Galluzzi et al., 2012). After 4 and 24 h exposure, cells were harvested by trypsinization, counted, and diluted in DMEM/F12 with 5% FBS, and 250 cells per well were seeded in 6-well plates (Corning^®^, USA). The cell medium was freshly replaced every 2 days. After 8 days, the formed cell colonies were fixed in PBS for 30 min using 3.4% v/v formaldehyde (4980.1, Carl Roth, Germany) and stained with 10% v/v of Giemsa solution (GS500, Sigma-Aldrich, USA) in Milli-Q water for 60 min. Representative images were taken using the Lionheart FX automated microscope (BioTek, USA), and the colonies were counted using ImageJ (Fiji, version 1.53c). The results are expressed as the percentage of colony forming efficiency (% CFE) according to equation 2 (Ponti et al., 2014).

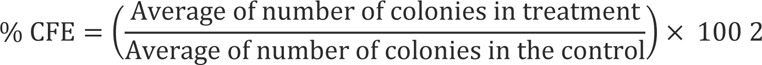

### DNA damage

To assess PM-induced DNA single- and double-strand breaks, the alkaline single-cell gel electrophoresis technique was performed according to the minigel comet assay method as demonstrated in a previous study (Di Bucchianico et al., 2017). After the exposure time, cells were detached using trypsin, and the cell suspension was diluted to a cell concentration of 2,500 cells mL^−1^. Then, each sample was mixed with 1% of low-melting-point agarose at 37°C, followed by embedding in a micro-agarose gel in microscope slides (2951-001, Thermo Fisher Scientific, USA). In our study, eight microgels per slide were arranged with untreated controls and PM-exposed cells, as well as a positive control, which was prepared by exposing cells to 30 µM hydrogen peroxide (H_2_O_2_) on ice for 5 min. Cells in the microgels were then lysed in a buffer (2.5 M NaCl, 0.1 M EDTA, 10 mM Tris, and 1% Triton X-100, pH 10) in the dark on ice for 1 h. To unwind the nuclear DNA, the slides were immersed in alkaline electrophoresis buffer (300 mM NaOH, 1 mM EDTA, pH >13) in the dark on ice for 30 min. Under an electric field generated by electrophoresis, the migration of damaged DNA toward the anode was achieved at 25 V with a current of 300 mA for 20 min. Then, each slide was rinsed with cold 0.4–M Tris (A411.1, Carl Roth, Germany) for 10 min, followed by rinsing in cold water for 10 min, and left for drying at least overnight. DNA was stained with 1:10,000 diluted SYBR Green (S11494, Thermo Fischer Scientific, USA) in TE (Tris–EDTA) buffer (10 mM Tris–HCl, 1 mM EDTA, pH 7.5). Images were taken using a Lionheart FX automated microscope to obtain micrographs at 20-fold magnification. One hundred nucleoids per minigel were analyzed using CometScore 2.0 software (TriTek Corp., USA), and the results are presented as mean % DNA in tail ± standard error of the mean of three independent exposures.

### Intracellular reactive oxygen species

The 2’,7’-dichlorodihydrofluorescein diacetate (H_2_DCF-DA) assay was used to investigate the potential of the PM to induce oxidative stress. For the intracellular ROS assay, A549 cells were seeded in 96-well plates at a density of 4 × 10^4^ cells cm^−2^ 24 h before exposure. The cells were preloaded with 100 µL of 20 µM H_2_DCF-DA in HBSS (with Ca^2+^ and Mg^2+^) for 30 min in a CO_2_ incubator at 37°C, followed by twice washing with prewarmed HBSS. The cells were then exposed to the PM suspension in DMEM (31053, Thermo Fisher Scientific, USA) without phenol red to avoid possible interference. As a positive control, 25 µM of tert-butyl hydroperoxide (8.14006, Sigma-Aldrich, USA) in HBSS was used. Immediately after the PM exposure, kinetic fluorescence was measured to quantify the increase of the fluorescence intensity with an excitation/emission wavelength of 485/535 nm in a microplate reader at 37°C (Varioskan^TM^ Lux, Thermo Fisher Scientific, USA) for 4 h. The detected ROS levels were expressed as the fold change of induced intracellular ROS.

### Hypoxanthine guanine phosphoribosyl transferase (HPRT) mammalian cell gene mutation

The *in vitro* hypoxanthine guanine phosphoribosyl transferase (HPRT) mutation assay was performed according to the OECD Guidelines for the Testing of Chemicals 476 (OECD476, 2016). Chinese hamster lung fibroblast cells V79-4 (CCL-3, ATCC^®^, USA) were freshly thawed and subcultured 2-4 times at a density of 2 × 10^5^ cells mL^−1^ in T-75 flasks (Corning^®^, USA). For PM exposure, 4 × 10^4^ cells were seeded in 12-well plates (Corning^®^, USA) and cultured for 24 h at 37°C with 5% CO_2_. After incubation, cells were exposed to a subcytotoxic concentration of PM (15 µg cm^−2^) in DMEM/F12 without serum for 24 h at 37°C with 5% CO_2_. The assay measures the disruption of HPRT enzyme activity in mutant cells after exposure to PM. It allows only the mutant cells to proliferate in the presence of purine analogue 6-thioguanine (6-TG) cytostatic effects, which inhibits the cellular metabolism of nonmutated cells.

For the experiment, cells were exposed to 15 µg cm^−2^ combustion PM of different fuel types for 24 h. At the same time, untreated cells were cultured in DMEM without FBS as a negative control, and cells were treated with 0.1 mM of ethyl-methanesulfonate (EMS, Sigma-Aldrich, USA) served as a positive control. After 24 h of exposure, 2 × 10^5^ cells were subcultured in T75 flasks (Corning^®^, USA) in a CO_2_ incubator at 37°C to allow the phenotypic expression of the induced mutations for an additional 11 days and split when confluent. On days 5 and 11, 100 cells per well were plated in 6-well plates for the concurrent CFE assay.

After the fixation of the mutations, 1 million cells were subcultured in 6-well plates to allow mutant cells to form colonies, and after 3 h, the cells were treated with 500 µg mL^−1^ of 6-TG to select mutant cells. After 8- to 9 days of cultivation in 6-TG-containing media, the mutant colonies were stained with Giemsa solution. The mutation frequency (MF) was calculated using equation 3 (Åkerlund et al., 2018).

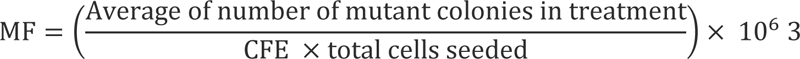

### CYP enzyme activation

To assess the bioaccessibility of PM-bound PAHs to cells, the induction of cytochrome P450 (CYP) activity was measured. As a suitable biomarker for the induced CYP, the 7-ethoxyresorufin-O-deethylase (EROD) was used to detect CYP1A1, CYP1B1, and CYP1A2 activities (Wen and Walle, 2005; Genies et al., 2013; Heinrich et al., 2014). In addition to EROD, 7-benzyloxyresorufin-O-debenzylase (BROD) was measured for detecting CYP1A1, CYP2B1, and CYP3A1 activities using the same principle (Hoet et al., 1997; Xie and Yang, 2018). After a 24 h PM exposure, cells were washed with prewarmed PBS and trypsinized for harvesting. As a positive control, cells were exposed to 5 µM of benzo[a]pyrene in DMEM/F12 without FBS for 1 h and harvested using the same procedure. The collected cells were centrifuged at 0.2 g for 5 min at room temperature, and the supernatant was removed. The dry pellets were stored at –80°C overnight for lysis until enzyme activity assay was performed. Cells were thawed, and the working solutions of the reaction mixtures were prepared from stock solutions of 10 mM of EROD (16122, Cayman, USA) and BROD (SC-208301, Santa Cruz, Germany) in dimethyl sulfoxide (DMSO). To prepare the EROD and BROD working solution, each 10 mM stock solution was further diluted to 10 µM in HBSS containing 0.1% v/v TritonX-100. After thawing the cells, 500 µL of each working solution was added to the cells, vortexed for 10 s, and incubated with CO_2_ at 37°C for 5 min. The initiation of the reaction was started by adding 250 µL of 1 mM nicotinamide adenine dinucleotide phosphate dispersed in Milli-Q water, followed by incubating at 37°C for 8 min to activate the enzymes (Hagemeyer et al., 2010). Then, 150 µL of the cell suspension was transferred to a black 96-well plate (SC-204450, Santa Cruz, Germany). Resorufin-associated fluorescence was quantified using a plate reader (Varioskan^TM^ Lux multimode microplate reader, Thermo Fisher Scientific, USA) with excitation and emission wavelengths set at 544 and 595 nm, respectively. The results of the induction of EROD and BROD activities were normalized with metabolic cell equivalents (MCE) derived from the trypan blue assay data of the corresponding treatment wells and expressed as pmol Resorufin × MCE^−1^ × min^−1^ (Heinrich et al., 2014).

### Interleukin-8 (IL-8) release

After 4 and 24 h of PM exposure, the culture media were collected, and the supernatants without cell debris and particles in the medium were separated via centrifugation at 110 g for 10 min (Universal 320R, Hettich, Germany) and stored at –80°C until analysis. The secretion of the pro-inflammatory IL-8 in the supernatant was determined in a 96-well plate using enzyme-linked immunosorbent assay according to the manufacturer’s instructions (DY208, R&D Systems Inc., USA). The assay used 3,3’,5,5’-tetramethylbenzidine as the substrate, and the absorbance of each sample was measured at 450 and 540 nm using a microplate reader (Varioskan™ Lux, Thermo Fisher Scientific, USA). The amount of measured proteins was calculated on the basis of standard curves and expressed as IL-8 pg mL^−1^.

### Statistical methods

All biological experiments were repeated 3 times and were analyzed using GraphPad Prism version 9.3.1 (GraphPad Software, USA) using one-way analysis of variance with the Dunnett’s test for multiple comparisons with *p*-values <0.05, <0.01, <0.001, <0.0001.

## Results and Discussion

### Characterization of emission particles from the research ship engine

The particle size distribution of different fuels was determined for particle diameters between 14 and 740 nm (Fig. S2). The results indicated that the particles from all HFOs showed remarkably skewed size distributions, with modes smaller than 100 nm, in contrast to the normally distributed MGO size distribution. This finding corroborates that of a previous study (Jeong et al., 2023) and is attributed to incomplete combustion of HFO fuel types and the formation of sulfuric acid particles, which depend on the inherent properties of marine fuel types (Zetterdahl et al., 2016; Lehtoranta et al., 2019). Interestingly, the particle size distribution of LS-HFO and HS-HFO_SYN_ showed a slightly bimodal distribution (Fig. S2), indicating interparticle phenomena in the exhaust line, such as particle-to-particle coagulation (Hinds, 2011).

The particle mass concentration of each fuel type is shown in Table 2. The combustion of HS-HFO_SYN_ produced the highest amount of particle mass, followed by HS-HFO, LS-HFO, ULS-HFO_AR_, and MGO. The particle mass concentration of MGO and ULS-HFO_AR_ was found to be in a lower range. Interestingly, the particle mass concentration of HS-HFO was between that of LS-HFO and HS-HFO_SYN_, indicating that parameters other than FSC played a role in the particle formation process. The measured eBC concentrations obtained from the aethalometer were found to be in accordance with the previously reported range (Jeong et al., 2023). The highest eBC mass concentration was observed for HS-HFO_AR_ emissions, followed by LS-HFO, HS-HFO, MGO, and ULS-HFO_AR_ (Table 2).

**Table 2.** Total particle and eBC mass concentrations from the exhaust emissions of each fuel type during the engine cycle in mg m^−3^ with standard deviations of three independent experiments (n = 3)

| [$\text{mg m}^{-3}$] | MGO | ULS-HFO<sub>AR</sub> | LS-HFO | HS-HFO<sub>SYN</sub> | HS-HFO |
| --- | --- | --- | --- | --- | --- |
| Mass concentration | $10 \pm 2$ | $12 \pm 1$ | $30 \pm 9$ | $79 \pm 4$ | $48 \pm 6$ |
| eBC concentration | $10 \pm 3$ | $7 \pm 1$ | $23 \pm 2$ | $43 \pm 2$ | $16 \pm 2$ |

### Physical characterization of PM in suspension

To define the physical characteristics of the suspended PM in the cell exposure medium, we used dynamic light scattering to analyze their hydrodynamic diameter, state of agglomeration, and stability. In general, the hydrodynamic diameters of PM were larger than their corresponding mobility diameters, with a mode of <100 nm (Fig. S2). This particle enlargement in the cell culture medium aligns with previous research, indicating that surface characteristics play a role in particle agglomeration and hydrodynamic dimeter increase (Kendall et al., 2015). Table 3 shows that HFO PM and MGO PM had hydrodynamic diameters of approximately 250–400 and approximately 200 nm, respectively.

**Table 3.** Results of DLS measurement in the exposure medium. All parameters are presented with standard deviations of three measurements.

|  | MGO | ULS-HFO<sub>AR</sub> | LS-HFO | HS-HFO<sub>SYN</sub> | HS-HFO |
| --- | --- | --- | --- | --- | --- |
|  | Exposure medium (DMEM/F12) |  |  |  |  |
| **Diameter [nm]** | 185 ± 2 | 310 ± 8 | 286 ± 4 | 398 ± 9 | 262 ± 8 |
| **PDI [-]** | 0.18 ± 0.01 | 0.3 ± 0.03 | 0.21 ± 0.01 | 0.34 ± 0.03 | 0.28 ± 0.02 |
| **Zeta-potential [mV]** | -18.6 ± 2.1 | -14.5 ± 1.0 | -13.6 ± 1.7 | -14.2 ± 1.0 | -11.9 ± 1.3 |

The PDI of the HFO PM samples was in the range of 0.2–0.35, indicating that they had a relatively broad size distribution in the exposure medium, whereas the MGO PM samples had a narrower size distribution with a PDI value below 0.2. To explore the potential stability of the colloidal system, we measured the zeta potential of the PM, which reflects the tendency for repulsion between particles due to their surface charge. The zeta-potential of MGO PM in the cell medium was found to be –18.6 mV, whereas that of HFO PM ranged from –12 mV to –14.5 mV.

### Chemical composition and mutagenic equivalent of the collected PM

The mass concentrations of 12 targeted PAHs are shown in Table 4 and Fig. S3. In general, PM-bound PAH concentrations for all HFOs were at least one order of magnitude higher than those for MGO, consistent with a previous study (Streibel et al., 2017).

**Table 4.**
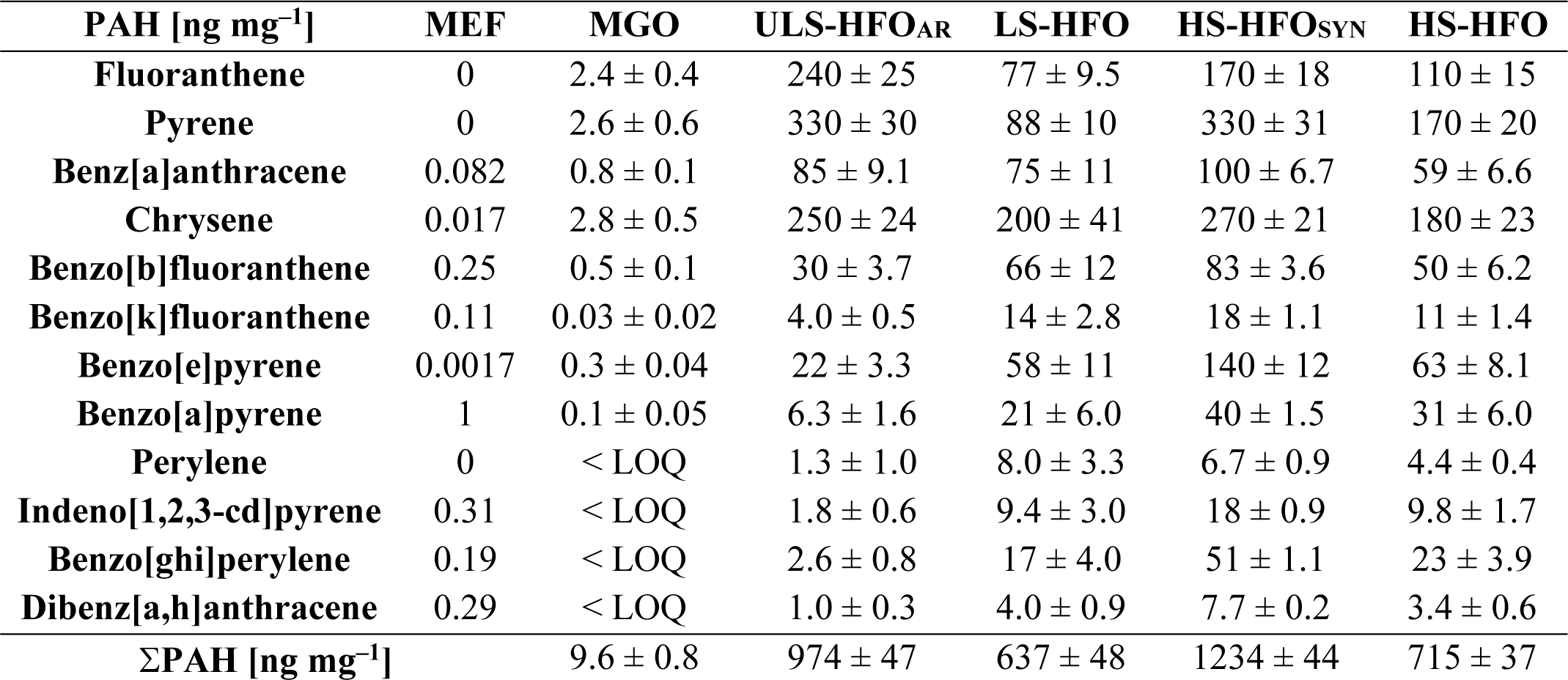
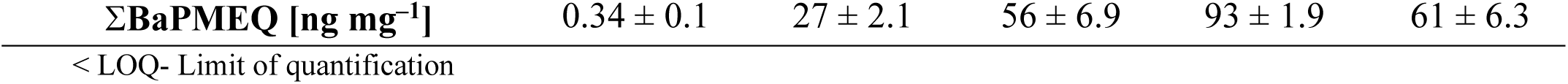
PM-bound PAHs, mutagenic equivalency factor (MEF) of 12 targeted PAHs (Durant et al., 1996) with standard deviations of three independent experiments (n = 3) and the sum of the mutagenic equivalent values (BaPMEQ) of the combustion PM from each fuel type.

Three dominant substances in the combustion particulate of all fuel types were pyrene, chrysene, and fluoranthene. We found a slightly lower relative contribution of fluoranthene and pyrene in this study compared with that in Sippula et al. (2014), which could be due to the much longer sampling time and associated losses of more volatile species due to blow-off effects in our study. The PM from ULS-HFO_AR_ showed an abundant emission profile of fluoranthene, pyrene, and benz[a]anthracene. The highest overall PAH concentrations were found for HS-HFO_SYN_ PM. In contrast, the fuel with the highest FSC, HS-HFO, showed average abundance for almost all PM-bound PAHs. The calculated MEQ values showed that the mutagenic potential of emitted PM from different marine fuel types differed significantly (Table 4). In general, the ΣBaPMEQ values of PM from all HFO fuel types were approximately 100 times higher than those of MGO PM. PM from HS-HFO_SYN_ had the highest total BaPMEQ of 93 ng mg^−1^, and >70% consisted of benz[a]anthracene, benzo[a]pyrene, benzo[b]fluoranthene (BbF), and benzo[ghi]perylene contributions. Other HFO PM also contained high concentrations of benzo[a]pyrene and benzo[b]fluoranthene. Interestingly, the PM from the aromatic-rich fuel ULS-HFO_AR_ showed the lowest total BaPMEQ value, mainly composed of 2- to 4-ring PAHs, especially alkylated phenanthrenes (Käfer et al., 2019). Despite having lower total PAH concentrations than ULS-HFO_AR_ PM, LS-HFO PM showed higher MEQ than ULS-HFO_AR_ PM due to the higher abundance of BaP and BbF as well as 5-ring PAHs such as indeno[1,2,3-cd]pyrene, benzo[ghi]perylene, and dibenz[a,h]anthracene, with higher MEQ compared with that of other PAHs. This trend aligns with the findings of a prior study conducted by Oeder et al. (2015), which also reported an increased concentration of these high molecular weight components. Although the MGO PM had a very low ΣBaPMEQ, it mainly consisted of BaP and BbF, which contributed to >60% of the ΣBaPMEQ of MGO PM. In contrast to other combustion sources, combustion aerosol particles from the use of both HFOs and diesel-like marine fuels can contain substantial amounts of alkylated derivatives of PAHs with three or more aromatic rings (Sippula et al., 2014; Czech et al., 2017). As these alkylated PAHs are not analyzed or listed with an appropriate MEF, the overall mutagenic potential of the combustion particles may be underestimated.

In addition to the PAH content, the elemental composition of the PM was analyzed using ICP-OES. As shown in Table 5, higher concentrations of vanadium, and sulfur per PM mass were detected with higher FSC, whereas zinc, and calcium concentrations in the exhaust showed no trend. Lube oil has been reported as a major source of zinc and calcium (Jung et al., 2003), and their ratio normalized to particle mass decreased with increasing FSC, whereas their absolute emissions did not change. Therefore, we observed significantly higher concentrations of zinc and calcium in the ULS-HFO_AR_ PM relative to PM mass. Iron and nickel concentrations did not show remarkable variations among the PM from different fuel types, except the PM from ULS-HFO_AR_, which had the lowest nickel concentration. Particulate emissions of nickel and vanadium are known to be mainly fuel derived and associated with the high-boiling fractions of crude oil but vary among different crude oil sources (Quimby et al., 1991). Therefore, the content of these metals in PM was reasonably high for the fuels with a high low-boiling constituents’ content. The elemental analysis of MGO PM was not possible because of the limited amount of collected PM. However, previous studies have indicated that the emission factors of transition metals (Fe, Ni, and V) from MGO combustion are significantly lower compared with those from HFO combustion (Table S3) (Sippula et al., 2014; Oeder et al., 2015; Hermansson et al., 2021; Sueur et al., 2023). The results of the elemental analysis normalized to PM mass are shown in Table S4.

**Table 5.** Elemental mass concentrations from the exhaust emission of each fuel type during the engine cycle expressed as µg mg^−1^ with the uncertainty of the instrumental analysis.

| **[$\mu\text{g mg}^{-1}$]** | **ULS-HFO<sub>AR</sub>** | **LS-HFO** | **HS-HFO<sub>SYN</sub>** | **HS-HFO** |
| --- | --- | --- | --- | --- |
| **Calcium (Ca)** | $45 \pm 0.4$ | $23 \pm 0.2$ | $10 \pm 0.1$ | $6.6 \pm 0.07$ |
| **Iron (Fe)** | $5.7 \pm 0.03$ | $4.3 \pm 0.03$ | $4.2 \pm 0.03$ | $5.1 \pm 0.03$ |
| **Nickel (Ni)** | $1.2 \pm 0.01$ | $6.2 \pm 0.08$ | $8.3 \pm 0.1$ | $5.9 \pm 0.07$ |
| **Sulfur (S)** | $29 \pm 0.2$ | $39 \pm 0.3$ | $28 \pm 0.2$ | $41 \pm 0.3$ |
| **Vanadium (V)** | $5.4 \pm 0.2$ | $14 \pm 0.4$ | $13 \pm 0.4$ | $24 \pm 0.7$ |
| **Zinc (Zn)** | $1.8 \pm 0.02$ | $0.7 \pm 0.01$ | $0.50 \pm 0.01$ | $0.20 \pm 0.01$ |

### Cell viability and clonogenicity

Cell viability was assessed using the trypan blue assay, which determines whether cells take up the dye due to cell membrane damage, after 4 and 24 h exposure, and the results are shown in Fig. 2a and 2b. Both 4 and 24 h exposure to the highest concentration of LS-HFO PM resulted in a significant reduction in cell viability, observing a decrease to 85%.

**Fig. 2.**
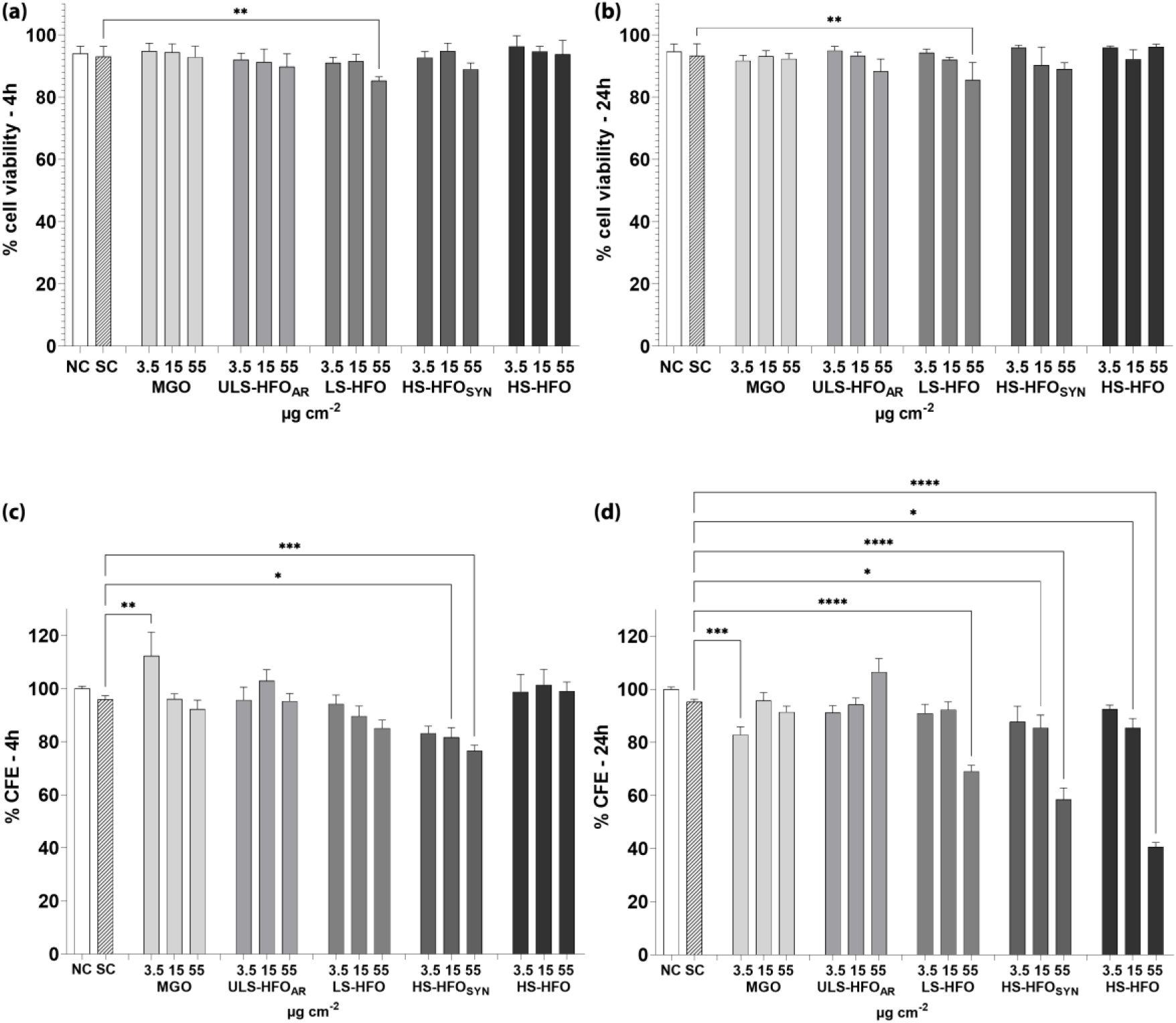
A549 cell viability and clonogenicity after exposure to different concentrations (3.5, 15, and 55 µg cm^−2^) of PM from marine fuel emissions. NC, negative control; SC, solvent control. Cell viability following (a) 4 and (b) 24 h exposure. % CFE following (c) 4 and (d) 24 h exposure Results are presented as mean ± SEM from three independent experiments (n = 3; \**p* < 0.05; \*\**p* < 0.01; \*\*\**p* < 0.001; \*\*\*\**p* < 0.0001 vs. SC

To investigate the effects of particles on cell clonogenicity, the CFE assay was performed and its relative colony forming efficiency (% CFE) after particle exposure is shown in Fig. 2c and 2d. In general, the CFE assay is known to be more sensitive than other cell viability assays such as trypan blue assay for detecting biological responses (Ponti et al., 2014), which was also observed in our experiment. HS-HFO_SYN_ PM caused large reductions in CFE at the intermediate (15 µg cm^−2^) and highest concentrations (Fig. 2c), likely due to relatively high concentrations of high molecular weight (HMW)-PAHs such as pyrene, benz[a]anthracene, benzo[e]pyrene and benzo[ghi]perylene. Interestingly, the MGO PM induced approximately 10% higher CFE after 4 h exposure at the lowest particle concentration (3.5 µg cm^−2^), whereas PM from the other fuel types did not cause significant alterations in CFE at this concentration (Fig. 2c).

Compared with the short exposure time, the long exposure time (24 h) revealed more pronounced cytotoxicity, especially for fuels with higher FSC (Fig. 2d). This remarkable reduction in CFE after 24 h may be due to increased PM uptake by cells and increased release of organic and inorganic compounds with prolonged exposure (Di Bucchianico et al., 2017; Corbin et al., 2018; Zerboni et al., 2019). CFE results showed that LS-HFO, HS-HFO_SYN_, and HS-HFO PM had significant cytotoxicity at approximately 70%, 60%, and 40%, respectively, at the highest concentration. Additionally, the intermediate concentration of both HS-HFO_SYN_ and HS-HFO PM reduced clonogenicity to approximately 85%. Comparable biological effects were reported in a PM exposure study on A549 cells, where HFO PM resulted in higher cytotoxicity and higher metabolic secretions, such as high lactic acid secretion compared with distillate oil PM (Oeder et al., 2015; Wu et al., 2018).

Contrary to the observed increase in CFE after 4 h exposure, MGO PM induced a significant reduction in CFE to 80% after 24 h at the same concentration (3.5 µg cm^−2^), whereas no reduction in CFE was observed for the other concentrations. This phenomenon may be related to the induction of fundamental metabolic functions caused by exposure to noncytotoxic concentrations of nanoparticles, which can result in an imbalance in adenine nucleotide homeostasis and mitochondrial malfunction (Fresta et al., 2018).

### Genomic instability, intracellular ROS, gene mutation, and IL-8 release in PM-exposed cells

PM-induced cellular responses have been associated with an imbalance of oxidants and antioxidants, which can lead to DNA damage, gene mutations, and chromosomal aberrations (Fresta et al., 2018; Cao et al., 2022; Cho et al., 2022). To address this phenomenon, the PM-induced DNA strand breaks and alkali-labile sites were examined and the results are presented as the percentage of DNA in tail after 4 and 24 h of particle exposure (Fig. 3a and 3b, respectively). The basal occurrence of DNA breaks was assessed in both the negative and solvent controls and was approximately 3% for both exposure times with no significant differences.

**Fig. 3.**
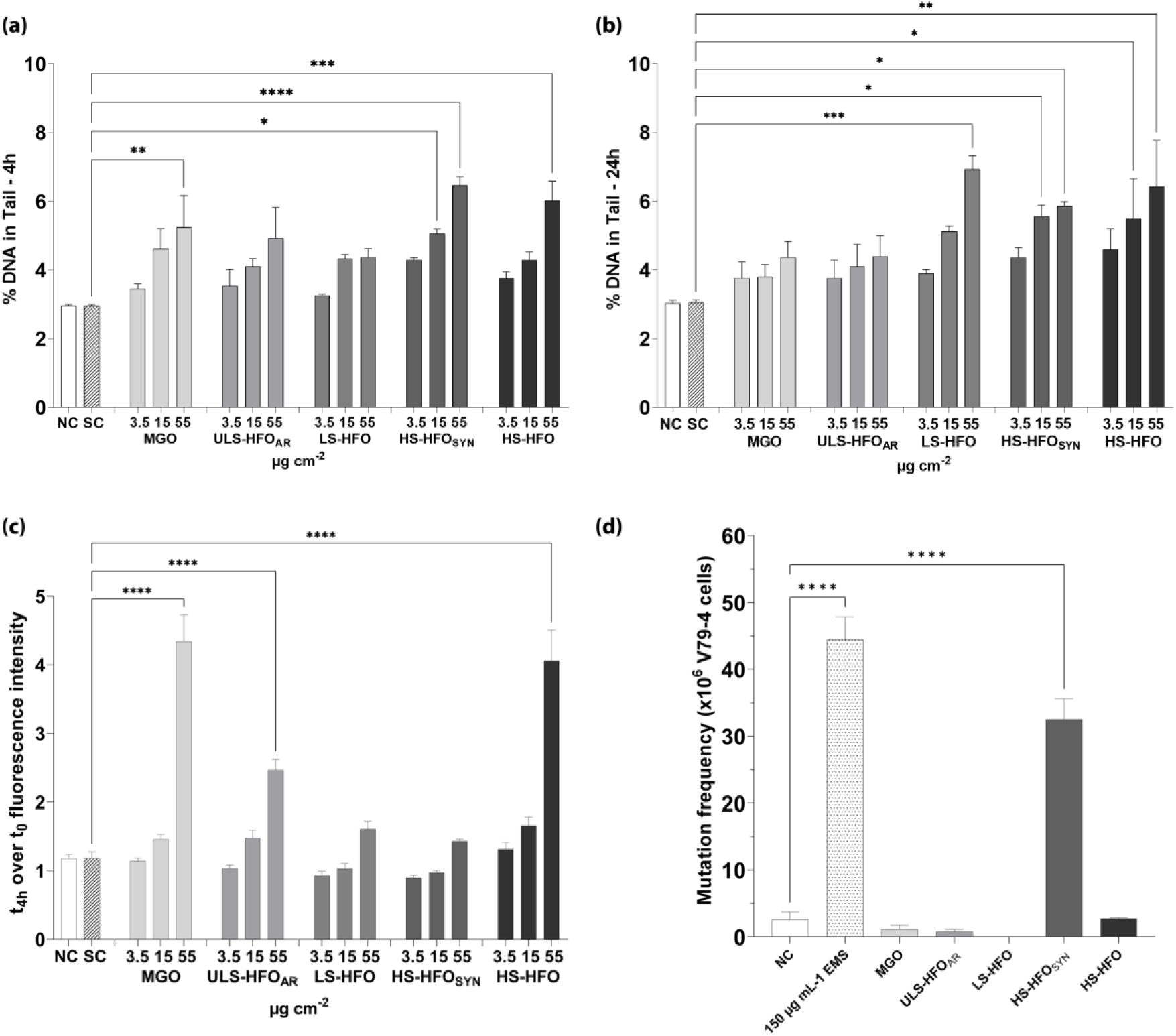
DNA damage, intracellular ROS, and V79-4 cell mutagenicity after exposure to different concentrations (3.5, 15, and 55 µg cm^−2^) of PM from marine fuel emissions. NC, negative control; SC, solvent control. DNA strand breaks measured using comet assay and represented as percentage of DNA damage in tail of A549 cells after 4 (a) and 24 h (b) exposure. (c) Intracellular ROS formation calculated as fold change of the fluorescence intensity kinetics in A549 cells over 4 h exposure. (d) HPRT gene mutation frequency in V79-4 cells after 24 h exposure to 15 µg cm^−2^ of marine fuel PM. Results are presented as mean ± SEM from three independent experiments (n = 3). \**p* < 0.05; \*\**p* < 0.01; \*\*\**p* < 0.001; \*\*\*\**p* < 0.0001 vs. SC (a, b, and c) or NC (d) for the mutation frequency

After 4 h exposure, MGO and ULS-HFO_AR_ PM increased the genotoxic effects with increasing concentration, but only MGO PM induced significant DNA damage at the highest concentration. Moreover, HS-HFO_SYN_ PM, one of the aromatic-rich fuels, induced significant DNA damage at the intermediate and highest concentrations. Similar DNA damage was observed at the highest concentration of HS-HFO PM (Fig. 3a). After 24 h exposure, LS-HFO PM caused significant DNA damage at the highest concentration. Furthermore, a slight reduction in DNA damage was observed for MGO PM at the intermediate and highest concentrations (Fig. 3b), which may indicate possible DNA repair mechanisms during a longer exposure time (Jugan et al., 2012).

To determine the role of PM-induced oxidative stress in the formation of DNA strand brakes, the production of intracellular ROS was investigated. The occurrence of intracellular ROS induced by PM exposure is demonstrated by a kinetic fluorescence intensity reading over 4-h exposure (Fig. 3c). An increase in intracellular ROS was observed with increasing concentrations of particles from all fuel types, significantly following exposure to MGO, ULS-HFO_AR_, and HS-HFO particles at the highest concentration with 4.3-, 2.5-, and 4-fold increases, respectively. Exposure to LS-HFO (1.6-fold) and HS-HFO_SYN_ (1.4-fold) PM showed a slight increase in ROS at the highest concentration, although it was not significant. MGO PM induced the highest intracellular ROS formation, which may be attributed to their relatively small size in the suspension compared with those of other PM (Table 3). These findings are consistent with that of previous studies that have shown that particle size can significantly influence their intracellular internalization, which in turn can affect ROS formation (Dos Santos et al., 2011).

In contrast, HS-HFO PM also induced a similar increase in ROS. Here, the high content of transition metal compounds in HS-HFO PM, especially vanadium (V[III]; (Popovicheva et al., 2009; Sueur et al., 2023), may play an important role in the impairment of mitochondrial functions and depletion of antioxidant glutathione (Leikauf et al., 2020; Pardo et al., 2020). Moreover, increased ROS formation can be caused indirectly by PM-bound organic compounds in a synergistic pathway (Dilger et al., 2016). Interestingly, our data showed a statistically significant ROS production in response to ULS-HFO_AR_ PM at the highest concentration despite its low eBC concentration, but higher iron content compared with that of MGO and HS-HFO PM. This phenomenon suggests intricate mechanisms for intracellular ROS production due to the complex mixtures of PM, which could only partially explain the observed primary DNA damage and its persistence.

To better assess the genotoxic and carcinogenic risks of exposure, we investigated the mutagenic potential of MGO and HFO PM, which could be directly and indirectly increased by the inadequate DNA repair and severe DNA breaks (Cho et al., 2022). The HPRT MF increased only after V79-4 cell exposure to HS-HFO_SYN_ PM, although LS-HFO and HS-HFO PM induced comparable DNA damage (Fig. 3d). These results aligned with the trend of the total BaPMEQ values, which were remarkably higher for HS-HFO PM than for other PM, such as from LS-HFO and HS-HFO, indicating the importance of PAH content in causing the observed mutagenic effects (Table 4).

Previous studies suggest that ROS generation and pro-oxidant cellular states are due to not only bioavailable PAHs but also their specific metabolic pathways (Briedé et al., 2004; Park et al., 2008). To gain insight into the metabolic processes of bioavailable PM-bound organic compounds, we measured the alteration of aryl hydrocarbon receptor activated phase I enzymes as an indicator of chemical surveillance (Lai et al., 2021).

Fig. 4a and 4b shows the enzymatic activity of xenobiotic metabolizing oxidoreductases of the CYP1A (EROD) and CYP2B (BROD) subfamilies after 24 h exposure. A slight induction of EROD by LS-HFO PM was observed with an increase in PM concentration, although not statistically significant. The PM from the aromatic-rich fuel types ULS-HFO_AR_, HS-HFO_SYN_, and HS-HFO induced EROD activity at the highest concentration. These results are consistent with those of a previous study showing significant activation of CYP1A1/1A2 enzymes induced by organic extracts of PAH-rich PM_10_ (Arrieta et al., 2003). The relatively high concentration of chrysene, abundant in ULS-HFO_AR_, and HS-HFO_SYN_ particles, may contribute to the significant induction of CYP1A1/1A2, as chrysene has been shown to induce significant activation of CYP1A1 and 1A2 in mouse liver cells (Shimada and Fujii-Kuriyama, 2004). Moreover, the presence of BaP and benz[a]anthracene is attributed to the increase of EROD activity, as shown in a previous *in vivo* study (Kim et al., 2003). The PM from aromatic-rich fuel types showed a nonsignificant, but still PM concentration-related induction of BROD. Only HS-HFO PM induced significant BROD activity at the highest exposure concentration (Fig. 4b), suggesting the presence of other chemical compounds not abundant in the PM from other fuel types. Neither EROD nor BROD was activated by LS-HFO PM, which contains relatively low PAH concentrations compared with other HFOs. The significant induction of EROD activity by ULS-HFO_AR_ PM may be due to the high levels of bioavailable low molecular weight (LMW)-PAHs contained in PM, which are more water-soluble than other PAHs (Tang et al., 2006). To further investigate this mechanism, we evaluated the occurrence of IL-8, one of the major pro-inflammatory factors, induced by CYP450 (Pei et al., 2002).

**Fig. 4.**
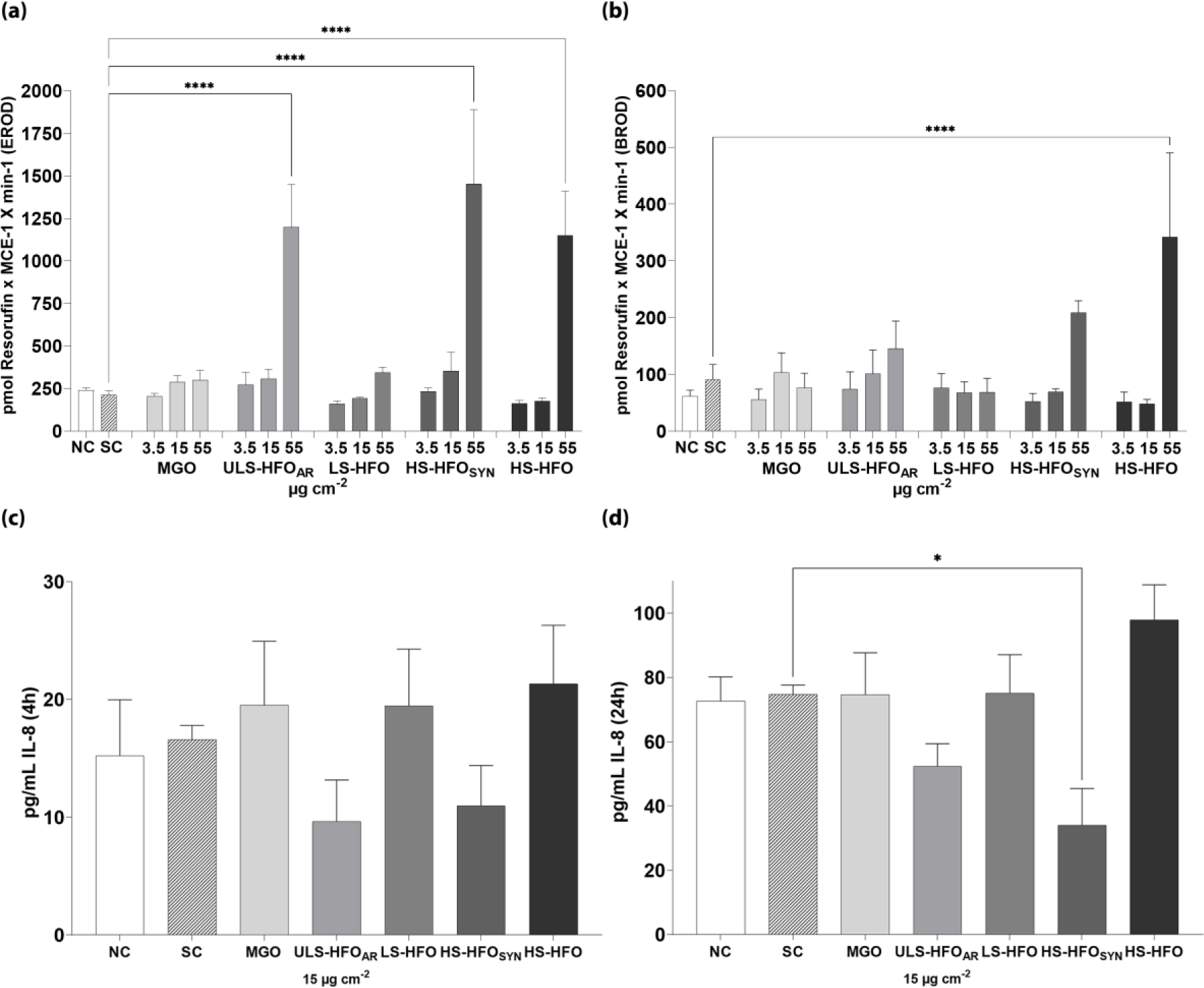
Induction of the Resorufin intensity from 7-ethoxyresorufin-O-deethylase (EROD) (a) and 7-benzyloxyresorufin-O-debenzylase (BROD) (b) in A549 cells normalized by metabolic cell equivalent (MCE) and exposure time to different concentrations (3.5, 15, and 55 µg cm^−2^) of PM from marine fuel emissions. The release of the pro-inflammatory cytokine IL-8 in A549 cells exposed to 15 µg cm^−2^ of marine fuel PM for 4 h (c) and 24 h (d). Data are shown as the mean + SE (n = 3). Statistically significant with respect to the solvent control according to one-way ANOVA with Dunnett’s test. \**p* < 0.05; \*\**p* < 0.01; \*\*\**p* < 0.001; \*\*\*\**p* < 0.0001 vs. the solvent control

Fig. 4c and 4d shows the release of IL-8 after both 4 and 24 h exposure to PM at a subcytotoxic particle concentration (15 µg cm^−2^). This PM concentration was chosen on the basis of the cytotoxicity results (trypan blue and CFE) where the cell viability was maintained above 80% so that the released IL-8 is not artificially miscalculated because of low cell viability. After 4 h of exposure, IL-8 release induced by PM from all fuel types did not show a statistically significant increase (Fig. 4c). However, HS-HFO_SYN_ PM caused significant suppression of IL-8 releases after 24 h exposure. This result may support our hypothesis that a disruption of signaling pathways may have occurred after the HS-HFO_SYN_ PM exposure. Additionally, a nonsignificant but slight suppression of IL-8 was caused by ULS-HFO_AR_ PM. The release of pro-inflammatory factor due to PM exposure resulted in a possible disruption of the cytoskeleton and trafficking of signal molecules, such as vesicles carrying IL-8 to the plasma membrane as shown by Longhin et al. (2018). These findings suggest a complex relationship between PM exposure-induced metabolic pathways and inflammatory responses. In contrast, HS-HFO PM induced a slight release of IL-8, which was not statistically significant, and may have contributed to the induction of intracellular ROS generation without inhibiting the transfer of signaling molecules. Furthermore, the results support the hypothesis of indirect ROS formation by PM-bound PAHs, as PM from aromatic-rich fuel types induced the activation of phase I enzymes regardless of their FSC.

## Conclusions

In this study, we investigated the effects of PM_2.5_ derived from ship emissions of different marine fuel types on A549 epithelial cells at two different exposure times. We evaluated their cytotoxicity, genotoxicity, intracellular ROS formation, activation of phase I enzymes, pro-inflammatory release, and potential to induce gene mutations in V79-4 cells. The marine fuels used in this study, including MGO and HFO with different FSCs, produced different particle emission profiles. Our research showed that fuel types with high levels of PAHs were more effective in inducing toxicological responses such as cytotoxicity, genotoxicity, and mutagenicity than distillate fuels such as MGO. Interestingly, we observed that these responses were not necessarily induced due to the FSC, but rather due to the inherent characteristics of PM, such as the content of PAH and size of the PM. It is worth noting that the biological responses induced by PM in our experiments were based on the same PM mass, as this study focused on the combustion particle quality related to the fuel type for environmental burden and health effects. Therefore, on the one hand, the generally higher formation of particle number and mass emission factors of HFOs than MGO must be considered when interpreting PM-driven toxicological effects. On the other hand, given the significant induction of intracellular ROS, the toxicological effects of MGO PM should also be taken into account, emphasizing the need for further reductions in particulate emissions. In addition, the relevance of alkylated PAHs to PM-induced toxicological responses can motivate further comprehensive research, as the number of possible isomers for alkylated PAHs increases with the number of added methylene units (Sun et al., 2014; Rüger et al., 2015). To enhance the relevance of such studies, we suggest considering aerosol exposure at the air liquid interface to mimic more realistic exposure scenarios and more efficient interactions between cells and particles than those in submerged conditions.

Overall, this study can contribute to a better understanding of the toxicological impacts of PM emitted by ships that use permitted marine fuel types with controlled FSC. Our study highlights the importance of considering the physical and chemical properties of marine fuel oils and their emissions in updating regulations. Therefore, it suggests that further legislation and restrictions of ship emissions concerning PM emission should be achieved using additional abatement systems to mitigate PM-related environmental and human health effects.

## Acknowledgement

We thank the Helmholtz Association of German Research Centers (HGF), Helmholtz International Lab *aeroHEALTH* (InterLabs-0005), University of Rostock and Bundeswehr University for supporting this project. We also thank Ms. Verena Häfner and Dr. Tobias Stöger (ILBD/CPC, Helmholtz Munich) for the active support regarding the physical characterization of particle suspensions. We gratefully acknowledge the Analytik Service AG (ASG) for the SIMDIS measurements and support.

## Author Contributions

Conceptualization: [Seongho Jeong, Thorsten Streibel, Martin Sklorz, Sebastiano Di Bucchianico, Ralf Zimmermann]; Methodology: [Seongho Jeong, Jürgen Schnelle-Kreis, Martin Sklorz, Sebastiano Di Bucchianico]; Formal analysis and investigation: [Seongho Jeong, Jana Pantzke, Svenja Offer, Uwe Käfer, Jan Bendl, Mohammad Saraji-Bozorgzad, Anja Huber, Bernhard Michalke, Uwe Etzien, Gert Jakobi, Jürgen Orasche, Hendryk Czech, Christopher P. Rüger, Martin Sklorz, Sebastiano Di Bucchianico]; Validation: [Seongho Jeong, Jana Pantzke, Svenja Offer, Uwe Käfer, Geri Jakobi, Jürgen Schnelle-Kreis, Martin Sklorz, Sebastiano Di Bucchianico]; Writing - original draft preparation: [Seongho Jeong, Jürgen Schnelle-Kreis, Martin Sklorz, Sebastiano Di Bucchianico]; Writing - review and editing: [Seongho Jeong, Jana Pantzke, Svenja Offer, Uwe Käfer, Hendryk Czech, Christopher P. Rüger, Thorsten Streibel, Martin Sklorz, Sebastiano Di Bucchianico, Ralf Zimmermann]; Funding acquisition: [Bert Buchholz, Thomas Adam, Ralf Zimmermann]; Resources: [Thorsten Streibel, Bert Buchholz, Thomas Adam, Ralf Zimmermann]; Supervision: [Martin Sklorz, Sebastiano Di Bucchianico, Ralf Zimmermann]

## Data Availability

The datasets generated during and/or analyzed during the current study available from the corresponding author on reasonable request.

## Declarations

### Ethics approval and consent to participate

Not applicable

### Consent for publication

Not applicable

## Abbreviations

AI: Aromaticity Index
BaP: Benzo[a]pyrene
BbF: Benzo[b]fluoranthene
BROD: 7- benzyloxyresorufin-O-debenzylase
BSA: Bovine Serum Albumin
CCAI: Calculated Carbon Aromaticity Index
CFE: Colony Forming Efficiency
CYP: Cytochrome P450
DF: Dilution Factor
DMEM: Dulbecco’s Modified Eagle Medium
DMSO: Dimethyl Sulfoxide
DNA: Deoxyribonucleic Acid
DTD-GC×GC-HRTOFMS: Direct Thermal Desorption comprehensive two-dimensional Gas Chromatography coupled to High-Resolution Time-Of-Flight Mass Spectrometry
eBC: Equivalent Black Carbon
EJD: Ejector Diluter
ELISA: Enzyme-Linked Immunosorbent Assay
EMS: Ethyl Methane Sulfonate
EROD: 7-ethoxyresorufin-O-deethylase
FBS: Fetal Bovine Serum
FSC: Fuel sulfur content
HBSS: Hank’s Balanced Salt Solution
HMW: High Molecular Weight
HFO: Heavy Fuel Oil
HPRT: Hypoxanthine-guanine phosphoribosyl transferase
ICP-OES: Inductively Coupled Plasma Atomic Emission Spectroscopy
IL-8: Interleukin-8
IMO: International Maritime Organization
LMW: Low Molecular Weight
MCE: Metabolic Cell Equivalent
MEF: Mutagenic Equivalency Factor
MEQ: Mutagenic Equivalent
MGO: Marine Gas Oil
PAH: Polycyclic Aromatic Hydrocarbon
PBS: Phosphate Buffered Saline
PDI: Poly Dispersity Index
PM: Particulate Matter
PTD: Porous tube diluter
ROS: Reactive Oxygen Species
SEM: Standard Error of Mean
SECA: Sulfur Emission Control Area
TE: Tris- EDTA;
TG: Thioguanine.

